# Phylodynamics of sunflower chlorotic mottle virus, a re-emerging pathosystem

**DOI:** 10.1101/829119

**Authors:** Dariel Cabrera Mederos, Carolina Torres, Nicolás Bejerman, Verónica Trucco, Sergio Lenardon, Michel Leiva Mora, Fabián Giolitti

## Abstract

Distribution and epidemiological patterns of sunflower chlorotic mottle virus (SCMoV) in sunflower (*Helianthus annuus* L.) growing areas in Argentina were studied from 2006 to 2017. The virus was detected exclusively in the Pampas region (Entre Ríos, Santa Fe, Córdoba, La Pampa and Buenos Aires provinces). Phylodynamic analyses performed using the coat protein gene of SCMoV isolates from sunflower and weeds dated the most recent common ancestor (MRCA) back to 1887 (HPD95% = 1572-1971), which coincides with the dates of sunflower introduction in Argentina. The MRCA was located in the south of Buenos Aires province and was associated with sunflower host (posterior probability for the ancestral host, ppah= 0.98). The Bayesian phylodynamic analyses revealed the dispersal patterns of SCMoV, suggesting a link between natural host diversity, crop displacement by human activities and virus spread.

## 1. Introduction

Sunflower (*Helianthus annuus* L.) is one of the most important oilseed crops in Argentina; indeed, more than 1 800 000 ha are planted annually, with the country ranking as one of the main producers and exporters of sunflower and byproducts worldwide (FAO, 2019). Several viruses naturally infecting sunflower have been reported and described worldwide (Gulya et al., 2002; Sharman et al., 2008; Giolitti et al., 2014; Cabrera Mederos et al., 2017). Sunflower chlorotic mottle virus (SCMoV) is considered a reemerging pathogen and is the most widely distributed of the potyviruses infecting cultivated and wild sunflower in Argentina (Lenardon et al., 2001). SCMoV was first observed in 1973 near Necochea, south of the Buenos Aires province, and was named necochense mosaic of sunflower (Kiehr-Delhey and Delhey, 1981).

SCMoV was detected in Paraná (Entre Ríos), Venado Tuerto (Santa Fe), Río Cuarto (Córdoba), Balcarce, Miramar and Necochea (Buenos Aires) using molecular methods (Lenardon, 1994; Lenardon et al., 2001). SCMoV was identified in five natural hosts: *H. annuus*, *H. petiolaris* L., *Eryngium* sp, *Dipsacus fullonum* L. and *Ibicella lutea* L. (Dujovny et al., 1998; Giolitti et al., 2009; Giolitti et al., 2012; Bejerman et al., 2013); to date, the complete genomic sequences of four SCMoV isolates have been reported (Bejerman et al., 2010; 2013). SCMoV is transmitted by mechanical inoculation and by aphids in a non-persistent manner, but it is not transmitted by seeds (Dujovny et al., 1998), limiting the virus ability to be dispersed over long distances. Moreover, *D. fullonum* is a biennial weed that plays an important role in SCMoV disease epidemiology, which serves as a potential inoculum source to virus infection in sunflower fields (Giolitti et al., 2009).

As most potyviruses, the biology of SCMoV has also been studied for several decades (Dujovny et al., 1998; Giolitti et al., 2009). However, the migration pathways as well as the evolutionary dynamics and timescale of this pathogen are still unknown. Thus, the aim of this study was to determine the geographical distribution and phylodynamics of SCMoV. The evolutionary rate of the capsid protein (CP) gene, the ancestral host and the spatiotemporal dynamics of SCMoV isolates were estimated. The results may be useful to gain knowledge of the virus epidemiology in order to develop sustainable management schemes.

## 2. Materials and methods

### 2.1 Geographical distribution of SCMoV

The main sunflower-cultivated areas of Argentina (Buenos Aires, La Pampa, Córdoba, Santa Fe, Entre Ríos, Chaco and Santiago del Estero provinces) were surveyed during the crop growing season. Samples of sunflower and weed leaves were collected from sunflower fields and nearby weeds from 2006 to 2017. To determine the SCMoV distribution, samples were lyophilized and stored at −20 °C until used. The presence of virus in symptomatic sunflower and weed plants was tested by enzyme-linked immunosorbent assay (ELISA), using specific antiserum to SCMoV (Dujovny et al., 1998) and confirmed by RT-PCR. For molecular detection of SCMoV, total RNA extraction was performed with RNeasy^®^ Plant Mini Kit (Qiagen, Germany), according to the manufacturer’s recommendations. First-strand cDNA synthesis was performed with 1 µg total RNA using Moloney murine leukemia virus (MMLV) reverse transcriptase enzyme (Promega, US), according to the manufacturer’s instructions, and M4T (5-GTTTTCCCAGTCACGAC(T)_15_-3) as the initial primer (Chen et al., 2001). PCR reactions were carried out using Kapa HiFi DNA polymerase enzyme (Kapa Biosystems, US) and specific primers: CP1: (5-GGTGACAACATAGATGCAGG-3) and CP2: (5-ACATGTTACGAACCCCAAGC-3), which allow the amplification of the entire CP gene of SCMoV (Giolitti et al., 2009). PCR reactions were performed using 4 μl of cDNA, 1 U of Kapa HiFi DNA polymerase enzyme, 300 μM dNTPs (each), and 0.3 μM of each primer in a final reaction volume of 50 μl. The PCR conditions included an initial incubation for 3 min at 95 °C, followed by 34 cycles of denaturation at 98 °C for 30 s, annealing at 55 °C for 45 s, extension for 90 s at 72 °C, and a final extension at 72 °C for 10 min.

### 2.2 Sequencing, recombination and phylogenetic analyses

The PCR products, obtained in duplicate from symptomatic plants, were purified using Wizard® SV Gel and PCR Clean-Up System (PROMEGA, US), and then directly sequenced in both directions by Macrogen Inc. The obtained sequences were assembled using the DNASTAR Lasergene SeqMan software (DNASTAR, Madison, WI) and a 780-bp consensus sequence was generated for each sample. The 30 CP gene sequences of SCMoV obtained in this work were deposited in GenBank (Accession number MG885770-99); details of these and previously reported sequences are shown in Suppl. Table 1. Sequence alignments were carried out using MUSCLE (Edgar, 2004) via SeaView software (Gouy et al., 2010). Pairwise identity analyses of the SCMoV sequences were carried out using the MUSCLE-based pairwise alignment option implemented in Sequence Demarcation Tool SDT v1.2 (Muhire et al., 2014).

Recombination analysis was done using Recombination Detection Program v. 4 (Martin et al., 2015). The potential recombination events were tested in the sequence alignments of CP genes using the RDP, GENECONV, BOOTSCAN, MAXIMUM CHI SQUARE, CHIMAERA, SISCAN and 3SEQ recombination detection methods. The probability of a putative recombination event was corrected by a Bonferroni procedure, with a cut-off of p = 0.01. Only recombination events that were detected by at least four methods and that had significant phylogenetic support (data not shown) were considered credible evidence of recombination. Those sequences with evidence of recombination were excluded from the subsequent analyses, according to Gibbs et al. (2017).

In addition, the degree of mutational saturation was tested using the index of substitution saturation (Iss) statistic in DAMBE software (Xia, 2017). The analysis was performed on all sites and on third codon positions. The Iss was significantly lower than the critical Iss value (Iss.c) (*P* < 0.0001), indicating that there is little saturation across the SCMoV sequences in our dataset and is adequate for phylogenetic reconstruction.

For phylogenetic reconstruction, the model of base substitution (GTR + I + G4) was estimated using the JMODELTEST v. 2.1.9 software (Darriba et al., 2012), according to the Akaike information criterion. The phylogenetic tree was obtained using Bayesian inference in MRBAYES v. 3.2.6 software (Ronquist et al., 2012). The analysis was run up to convergence, which was assessed by effective sample size values higher than 200 using the TRACER v. 1.7.1 (Rambaut et al., 2018), and the first 10% of generations were discarded as burn-in.

### 2.3 Phylodynamic analysis

The non-recombinant SCMoV sequences of the CP gene obtained in this work and other four sequences available in GenBank (GU181199, GU181200, JN863232, JN863233) were subjected to a Bayesian coalescent analysis to study the spatiotemporal dynamics of the virus. To evaluate the temporal structure of the data, date-randomization test was performed with 20 replicas, using the TipDatingBeast R package (Rieux and Khatchikian, 2017). This analysis showed overlapping in the posterior probability distributions of the time to the most recent common ancestor (tMRCA) and the mean substitution rate estimated with the real data and the replica, indicating an insufficient temporal structure of the dataset using sampling times. Then, to calibrate the analyses, an external substitution rate of 1.15 × 10^−4^ nucleotide substitutions/site/year (s/s/y) was used, which was previously estimated for the CP gene of potyviruses (Gibbs et al., 2008).

Analyses were run using the substitution model (GTR + I + G4) selected according to the Akaike information criterion, estimated with JMODELTEST v. 2.1.9 software (Darriba et al., 2012), the uncorrelated lognormal molecular clock model (UCLN), and the GMRF Bayesian Skyride coalescent model (Minin et al., 2008). A spatial diffusion process was modeled along time-measured genealogies over continuous sampling locations (geographical coordinates) using different diffusion models (Homogenous Gamma, Cauchy and LogNormal distributions) (Lemey et al., 2010), implemented in the BEAST v. 1.10.4 software package (Suchard et al., 2018). In addition, since the sequences were isolated from different hosts (Suppl. Table 1), an asymmetric discrete trait model was used to estimate the ancestral hosts and their probability (Lemey et al., 2009). Convergence was assessed by effective sample size (ESS) values higher than 200, using Tracer v. 1.7.1 (Rambaut et al., 2018), and 10% of the sampling was discarded as burn-in.

Analyses with different diffusion models were compared via the Bayes factor using the marginal likelihood values estimated by stepping stone (SS) method (Baele et al., 2012). The model selected was the Gamma distribution (Suppl. Table 2). Two runs were combined and the most clade credibility tree (MCCT) was obtained. Uncertainty in parameter estimates was evaluated in the 95% highest posterior density (HPD95%) interval for most parameters or in the HPD80% for locations and diffusion rate. Visualization file of the spatiotemporal dynamic process was built using the SPREAD software (Bielejec et al., 2011).

## 3. Results

### 3.1 Geographical distribution of SCMoV

Leaves of sunflower and weeds, which exhibited the SCMoV-like symptoms (Suppl. Fig. 1), were positive by ELISA. In addition, the presence of SCMoV was confirmed by the amplification of a 780-bp fragment by RT-PCR. SCMoV was detected in 19 departments located in Entre Ríos, Santa Fe, Córdoba, La Pampa and Buenos Aires provinces. In sunflower fields from Chaco and Santiago del Estero provinces the disease was not detected (data not shown). Numerical values were assigned to locations where the disease was detected to show the geographical distribution (Fig. 1; Suppl. Table 1).

**Fig. 1.**
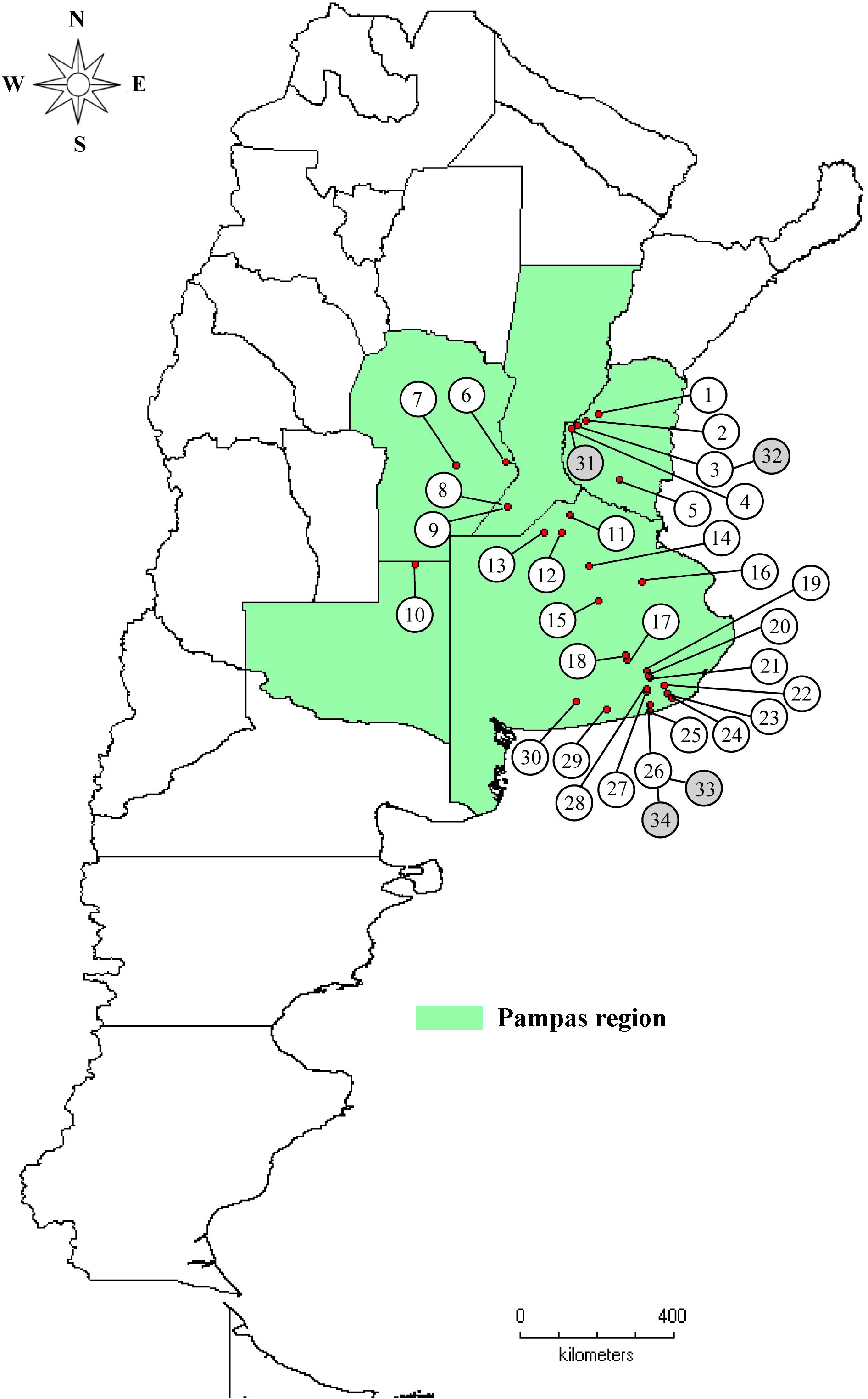
Geographical distribution of sunflower chlorotic mottle virus in Argentina. Locations where viral disease was detected are numbered from 1 to 34, and are shown in Suppl. Table 1.

### 3.2 Phylogenetic and recombination analyses

The pairwise analysis of the SCMoV isolates (n=34) showed nucleotide sequence identity values that ranged from 87.0 to 99.8%; these values did not show any association with the host, year of collection or geographical location. The highest variability was observed among isolates from Buenos Aires province: Pieres (GU181200, JN863233), Tandil (MG885780), Mechongue (MG885777), Miramar (MG885776) and Lobería (MG885799) (Suppl. Fig. 2). The pairwise analysis of the deduced amino acid sequences of the CP showed identity values that ranged from 94.2 to 100% (data not shown). A potential recombination event was detected within the CP gene of the analyzed SCMoV isolates, with the General Alvear isolate being the recombinant sequence, whereas Los Pinos and Pieres isolates would be the parental sequences (Suppl. Table 3). Since recombination confounds phylogenetic analysis (Gibbs et al., 2017), the recombinant sequence (MG885794) was discarded from the dataset.

In the phylogenetic tree, four main groups with high support values were observed, named as Groups 1 to 4 (Fig. 2), whereas it was not possible to classify two sequences with confidence (MG885795, MG885789). Interestingly, the more numerous groups (1 and 2) did not segregate according to geographical origin or their host. Group 1 included sequences from all the evaluated provinces, group 2 included sequences from Buenos Aires, Entre Ríos and Córdoba provinces and groups 3 and 4 included only sequences from Buenos Aires province (Fig. 2).

**Fig. 2.**
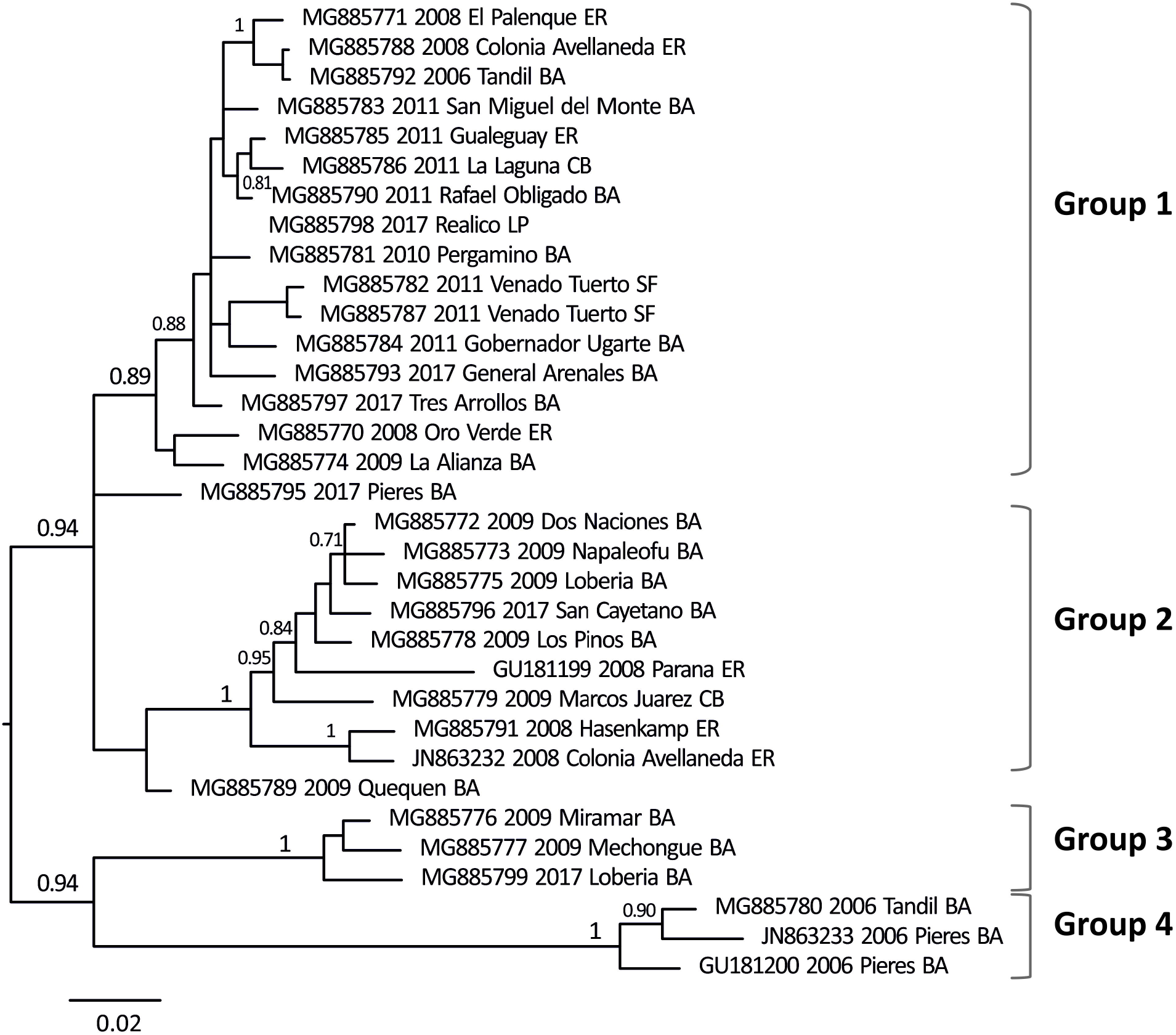
Majority rule consensus tree from a Bayesian analysis of sunflower chlorotic mottle virus CP gene sequences (780 nt). The tree was rooted according to the position estimated in the coalescent analysis. Posterior probability values higher than 0.7 are shown at nodes for relevant groups. Year of collection, location and province (abbreviated in uppercase) are indicated next to the accession number of the isolates. ER: Entre Ríos, CB: Córdoba, SF: Santa Fe, LP: La Pampa, BA: Buenos Aires.

### 3.3 Phylodynamic analysis

The most recent common ancestor for the SCMoV was dated back to 1887 (HPD95%= 1572-1971), and was located in the south of Buenos Aires Province (Lobería) (Fig. 3; Fig. 4 and Supplementary file), which would have associated with sunflower as host (posterior probability for the ancestral host, ppah= 0.98) (Fig. 3). This ancestral virus diversified in two clusters in ~1910-1920 that originated the four groups described above, all associated with sunflower as ancestral host (Fig. 3). Groups 1 and 2 diversified in 1943 (HPD95%= 1760-1990) and 1944 (HPD95%= 1760-1988), respectively, and would represent the most dispersed groups, including sequences from different locations. In particular, a transition from sunflower to *D. fullonum* was estimated in group 1 lineages between 1958-1967; since then, lineages cocirculated in both hosts. Group 3 lineages would have diversified in 1977 (HPD95%= 1874-2003) and still circulate in the present, whereas Group 4 lineages would have diversified in 1979 (HPD95%= 1860-1998) and showed an internal transition from sunflower to *D. fullonum* in 1985.

**Fig. 3.**
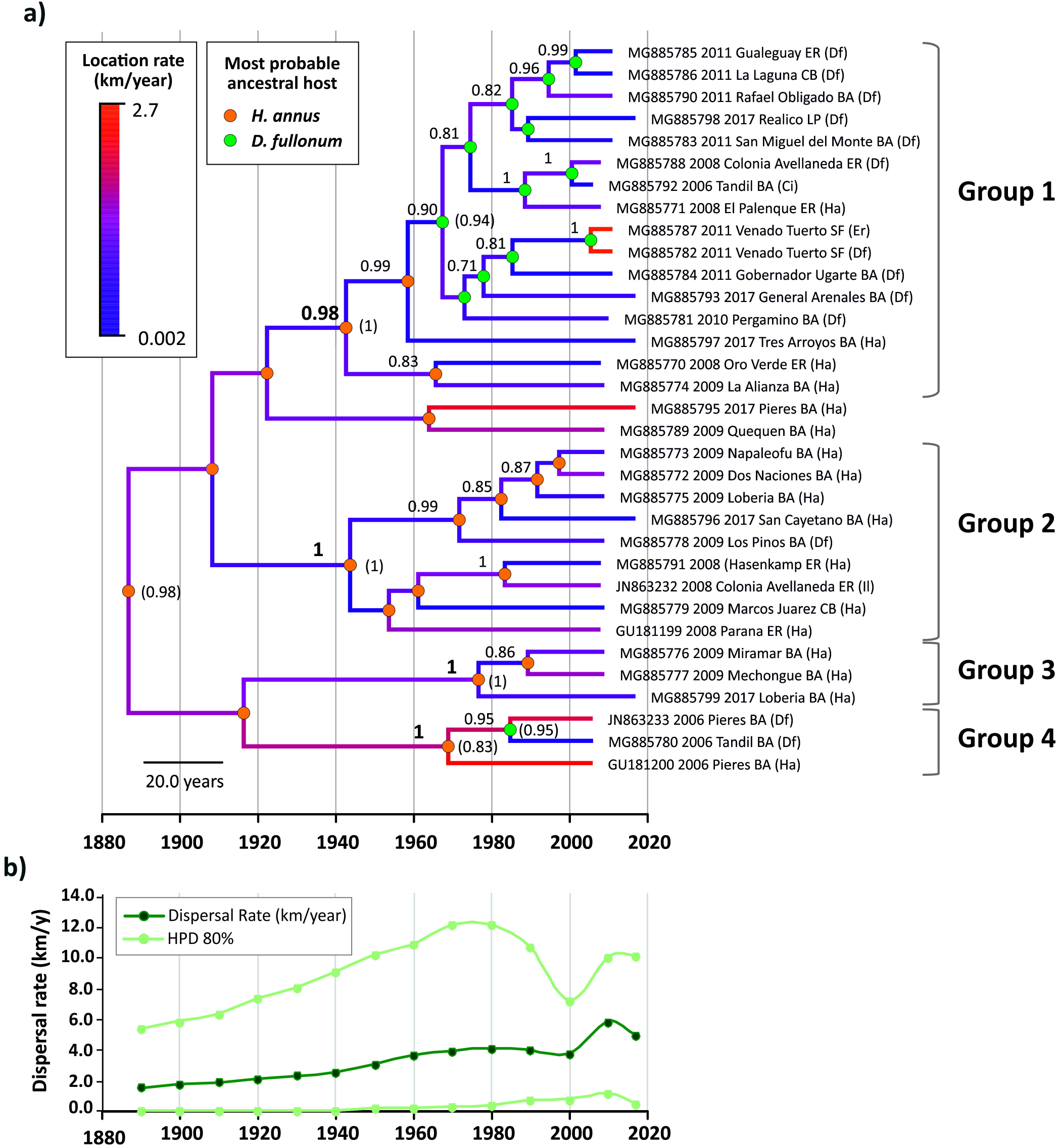
Maximum clade credibility tree of the posterior distribution of the Bayesian coalescent analysis of sunflower chlorotic mottle virus CP gene sequences (a) and a plot of the estimated dispersal rate over time (b). The tree is in temporal scale (years) and clade posterior probability values higher than 0.7 are shown at nodes. The blue-red color gradient of the branches represents relative dispersal rates (slower-faster) and the color at nodes represent the most probable ancestral host (posterior probability values for the ancestral host are shown in parenthesis, for relevant groups). Year of collection, location, province and host (abbreviated in uppercase) are indicated after the accession number of the isolates. Ha: *Helianthus annuus*, Df: *Dipsacus fullonum*, Ci: *Ciconium* sp., Er: *Eryngium* sp., Il: *Ibicella lutea*.

**Fig. 4.**
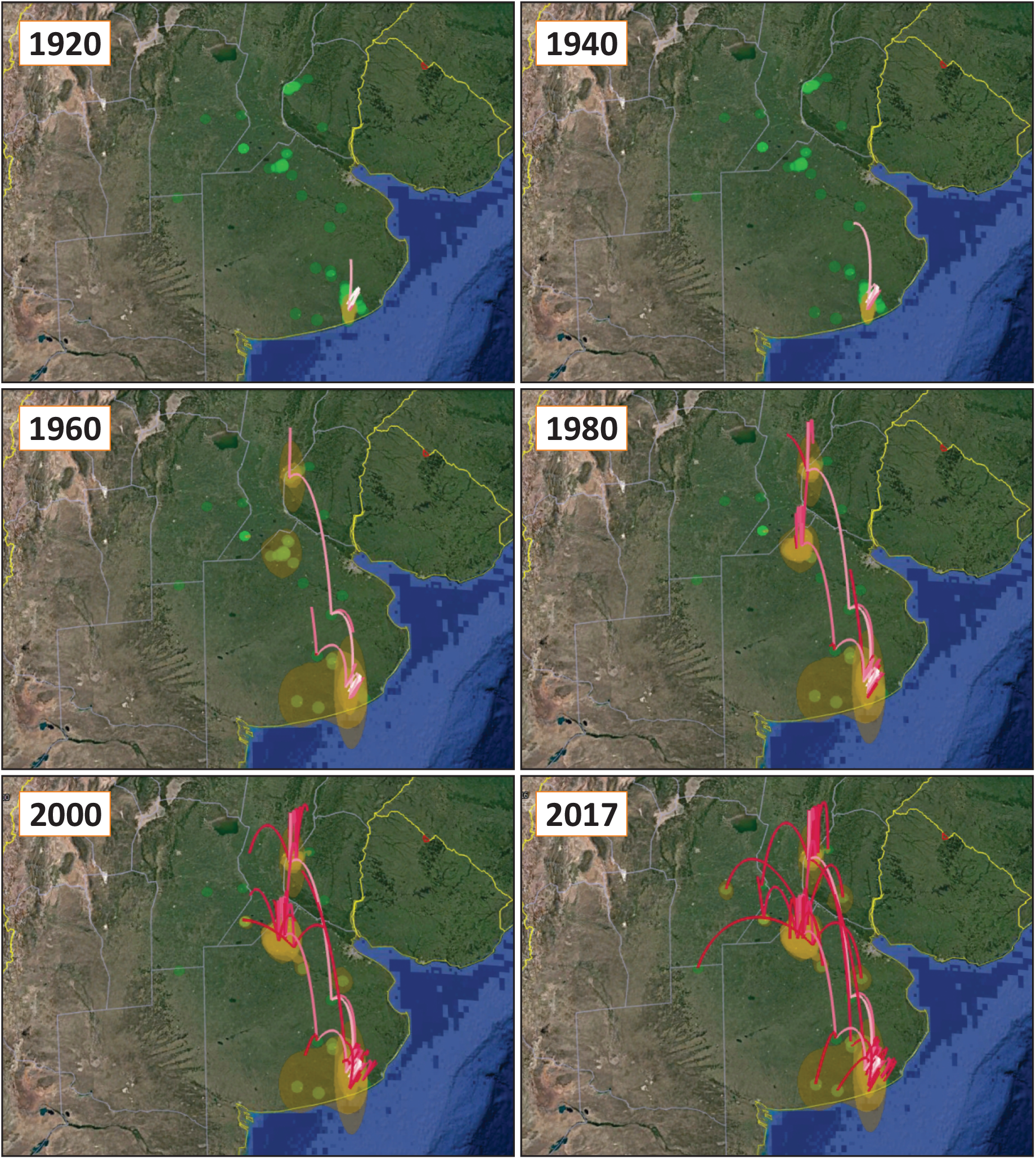
Spatiotemporal diffusion processes for the sunflower chlorotic mottle virus. Lines correspond to the Maximum Clade Credibility Tree projected on the surface, where white-red color gradient represents the relative age of the dispersal pattern (older-recent). Sampling sites are shown as green circles and the uncertainty on the estimated locations is represented by orange polygons (HPD80%).

The median dispersal rate was estimated as 4.0 km/year (HPD95%=0.6 – 8.2), showing a slight increase from the past to the present (Fig. 3). On the other hand, although an external substitution rate of 1.15 × 10^−4^ s/s/y was used as prior to calibrate the coalescent analyses, the posterior distribution of its estimate resulted in 5.1 × 10^−4^ s/s/y (HPD95%= 8.2 × 10^−5^ – 1.0 × 10^−3^). In addition, the demographic reconstruction did not show significant changes in the number of infections over time (data not shown).

## 4. Discussion

In the surveys conducted in the main sunflower producing regions of Argentina from 2006 to 2017, we confirmed the wide distribution of SCMoV; however, this virus was not detected in all the main producing regions. Although important sunflower areas are also located in Chaco region (Chaco and Santiago del Estero provinces), SCMoV was detected exclusively in the Pampas region of Argentina (Entre Ríos, Santa Fe, Córdoba, La Pampa and Buenos Aires provinces). In addition to the SCMoV hosts previously reported (Giolitti et al., 2009; 2012), in this work a new host was detected (*Ciconium* sp.). Moreover, the detection of SCMoV only in the Pampas region could be associated with the distribution not only of sunflower but also to the distribution of its natural host *D. fullonum* in Argentina (Daddario et al., 2015). This weed species was introduced from Eurasia and naturalized in Argentina (Burkart, 1957). It is a biennial species that reproduces only by seed (Busso et al., 2013), which are dispersed by water, animals and humans. *D. fullonum* seed germination near sunflower fields allows virus survival during periods when the main crop is absent. For this reason, *D. fullonum* plays an important role in SCMoV disease epidemiology.

Phylogenetic analysis of SCMoV revealed major diversity among isolates from Buenos Aires province, which coincided with the possible diversification center of this virus. Moreover, it was not possible to determine the association between genetic diversity and geographical localization or host, as previously observed for other potyviruses (Cabrera Mederos et al., 2019). In the Bayesian analysis, no structure of sequences from group 1 and 2 in relation to the geographical distribution was observed, indicating dispersion of the virus among locations over time. However, an apparent structure was observed in the isolates from groups 3 and 4, which provides strong evidence for the in situ evolution of viruses (Simmons et al., 2008).

The results of the phylogenetic analysis suggest that SCMoV can be subjected to dispersion to different regions, perhaps associated with the presence of infected hosts and mediated by the local spread by aphids in nearby areas. SCMoV has not been found to be transmitted by seeds (Dujovny et al., 1998; Giolitti et al., 2009); which clearly reduces the virus spread over long distances and could explain the observed dispersion rates (4.0 km/year). Interestingly, the dispersal rate doubled its value from 1930 to 1980, coinciding with the sunflower crop expansion in Argentina (Castaño, 2018). Moreover, between 2000 and 2010, an increment of SCMoV dispersal rate was observed; however, the sunflower-cropped area decreased compared with the period from 1995 to 2000 (ASAGIR, 2017; Castaño, 2018). The crop displacement associated with the production of soybeans and barley in Argentina, as well as the cultivation of sunflower in areas that have not been cultivated with this crop for a long period have also influenced the SCMoV spread. These scenarios also promote new encounters among virus-vectors and native or naturalized host, in which the host jump may occur (Jones, 2009), as observed in the transitions from sunflower to *D. fullonum*.

In the present analysis the ancestral origin of SCMoV diversification was estimated in the south of Buenos Aires province approximately 130 years ago. Moreover, the ancestral host was sunflower, which revealed a correlation with sunflower introduction in Argentina (Castaño, 2018). Other evidences also suggests that SCMoV originated in Argentina. For example, sunflower came originally from the United States (US) and was introduced to Europe toward the middle of the 16th century (Patiño, 1963). Interestingly, this virus has not been detected in US or European countries. Towards the end of the 19th century, with the Jewish immigration from Russia to Argentina, the first colonies established in the provinces of Santa Fe, Entre Ríos, Chaco and Buenos Aires made the first introductions of sunflower seeds. Moreover, the first important sunflower plantations were established in the department of Carlos Casares, Buenos Aires province (Castaño, 2018). Sunflower crops were multiplied and distributed in different regions of Argentina between 1881 and 1900. The results obtained in the phylodynamic analysis are consistent with the periods of sunflower dispersion in Argentina. Moreover, the transmission mode of SCMoV is the limiting factor for virus dispersion between distant regions, as observed in this study.

In previous studies, Giolitti et al. (2010) showed that SCMoV might be a pioneer species descending from PVY, which acquired infectivity to other plant families. Interestingly, the origins and/or first records of some potyviruses, including members of the PVY group, were in South America (Gibbs and Ohshima, 2010; Gibbs et al., 2017). PVY is a major pathogen of potatoes and other solanaceous crops worldwide that infects various plant species, including *Lactuca serriola*, a species of the *Asteraceae* family (Kaliciak and Syller, 2009). However, Dujovny et al. (1998) analyzed the host range of SCMoV means of mechanical inoculation, and detected the infection only in *Nicotiana occidentalis* and *Datura metel* among the tested *Solanaceae* species. Moreover, the *Asteraceae* species *Bidens subalternans* D.C., *Chrysanthemum coronarium* L., *Tagetes erecta* L., and *Zinnia elegans* Jacq were also infected experimentally, raising new questions: is SCMoV naturally transmitted in these weed species?, can SCMoV be transmitted by seeds in weeds species? Supposing that the ancestors of the present SCMoV populations depended on natural host diversity and aphid vector populations for their maintenance and spread, then the estimated MRCAs are restricted to natural dispersion in sunflower between localizations, in contrast to the situation with other virus dispersed by infected seed and other agricultural practices (Yasaka et al., 2017). This aspect could be related to the speciation and evolution of different isolates, transmission modes and natural host diversity. These findings provide epidemiological evidence about phylodynamic patterns of SCMoV, such as the knowledge about novel host species, diversification center, ancestral host and origin, which could be used in virus evolution studies and in implementation of control strategies.

## Acknowledgements

The authors would like to thank the Consejo Nacional de Investigaciones Científicas y Técnicas (CONICET) (Postdoctoral fellowship internal PDTS-Resolution No. 4885 and INTA-CIAP-IPAVE for financial support.

## Ethical approval

This article does not contain any studies with human participants or animals performed by any of the author.

## Appendix A. Supplementary material

**Suppl. Fig. 1.** Sunflower chlorotic mottle virus symptoms (leaf mosaic, mottling, yellowing patches) induced in naturally infected sunflower crops and weeds in Argentina. A: *Helianthus annuus*, B: *Dipsacus fullonum*, C: *Ciconium* sp., D: *Eryngium* sp., E: Ibicella lutea.

**Suppl. Fig. 2.** Pairwise analysis among nucleotide sequences of the coat protein gen of sunflower chlorotic mottle virus isolates from Argentina. Pairwise sequence identities were calculated using SDT v1.2 (Muhire et al. 2014) program.

**Supplementary file.** Spatiotemporal diffusion processes for the sunflower chlorotic mottle virus (in Keyhole Markup Language format that can be viewed using Google Earth or any other software capable of reading this format). See caption of Fig. 4 for details.

### Supplementary material

**Suppl. Table 1:** Geographical localization, collection date, host an accessions of sunflower chlorotic mottle virus isolates infecting sunflower and weeds in Argentina.

**Suppl. Table 2:** Bayes Factors (BF) for comparison of different diffusion models with marginal likelihood estimators calculated using Stepping Stone (SS) method.

**Suppl. Table 3:** Recombination sites in the coat protein gene of sunflower chlorotic mottle virus isolate detected by more of four recombination-detecting programs implemented in RDP4 software.

## References

ASAGIR. 2017. Historia. El girasol en la Argentina. Available from http://www.asagir.org.ar/acerca-de-historia-456 (2019/09/8).

Baele, G., Lemey, P., Bedford, T., Rambaut, A., Suchard, M.A., Alekseyenko, A.V., 2012. Improving the accuracy of demographic and molecular clock model comparison while accommodating phylogenetic uncertainty. Mol. Biol. Evol. 29, 2157–2167.

Burkart, A., 1957. Las Dipsacaceas asilvestradas en la Argentina. Bol. Soc. Arg. Bot. 6: 243–247.

Busso, C.A., Bentivegna, D.J., Fernández, O.A., 2013. A review on invasive plants in rangelands of Argentina. Interciencia 38, 95–103.

Bejerman, N., Giolitti, F., de Breuil, S., Lenardon, S. 2010. Molecular characterization of Sunflower chlorotic mottle virus: a member of a distinct species in the genus Potyvirus. Arch. Virol. 155, 1331–1335.

Bejerman, N., Giolitti, F., de Breuil, S., Lenardon, S., 2013. Sequencing of two sunflower chlorotic mottle virus isolates obtained from different natural hosts shed light on its evolutionary history. Virus Genes 46, 105–110.

Bielejec, F., Rambaut, A., Suchard, M.A., Lemey, P., 2011. SPREAD: spatial phylogenetic reconstruction of evolutionary dynamics. Bioinformatics 27, 2910–2912.

Cabrera Mederos, D., Bejerman, N., Trucco, V., de Breuil, S., Lenardon, S., Giolitti, F. 2017. Complete genome sequence of sunflower ring blotch virus, a new potyvirus infecting sunflower in Argentina. Arch. Virol. 162, 1787–1790.

Cabrera Mederos, D., Giolitti, F., Torres, C., Portal, O., 2019. Distribution and phylodynamics of papaya ringspot virus on Carica papaya in Cuba. Plant Pathol. 68, 239–250.

Castaño, F.D., 2018. The sunflower crop in Argentina: past, present and potential future. OCL-Oilseeds and fats, Crops and Lipids, 25, D105.

Chen, J., Adams M.J., 2001. A universal PCR primer to detect members of the Potyviridae and its use to examine the taxonomic status of several members of the family. Arch. Virol. 146, 757–766.

Daddario, J.F., Tucat G., Bentivegna, D., Anderson, F.E., 2015. La “carda silvestre” en Argentina: antecedentes, problemática causada por la invasión y estudios locales para resolverla. Boletín electrónico CERZOS, CONICET.

Darriba, D., Taboada, G.L., Doallo, R., Posada, D., 2012. jModelTest 2: more models, new heuristics and parallel computing. Nat. Methods 9, 772.

Dujovny, G., Usugi T., Shohara K., Lenardon S., 1998. Characterization of a potyvirus infecting sunflower in Argentina. Plant Dis. 82, 470–474.

Edgar, R.C., 2004. MUSCLE: multiple sequence alignment with high accuracy and high throughput. Nucleic Acids Res. 32, 1792–1797.

FAO., 2019. FAO Statistics. Available online: http://www.faostat.org.

Giolitti, F., Bejerman N., Trucco V., de Breuil S., Lenardon S., 2012. Caraguatá” (*Eryngium* sp.): Nuevo reservorio natural del sunflower chlorotic mottle virus (SCMoV). pp. 40. XIV Jornadas fitosanitarias Argentinas. 3-5/10, Potrero de los Funes, San Luis, Argentina.

Giolitti, F., Bejerman, N., Lenardon, S., 2009. *Dipsacus fullonum*: an alternative host of sunflower chlorotic mottle virus in Argentina. J. Phytopathol. 157, 325–328.

Giolitti, F., Bejerman N., Nome, C., Visintin, G., de Breuil, S., Lenardon, S. 2014. Biological and molecular characterization of an isolate of pelargonium zonate spot virus in infecting sunflower in Argentina. J. Plant Pathol. 96:189–194.

Gibbs, A.J., Ohshima, K., 2010. Potyviruses and the digital revolution. Annu. Rev. Phytopathol. 48, 205–223.

Gibbs, A.J., Ohshima, K., Phillips, M.J., Gibbs, M.J., 2008. The prehistory of potyviruses: their initial radiation was during the dawn of agriculture. PLoS one 3, e2523.

Gibbs, A.J., Ohshima, K., Yasaka R., Mohammadi M., Gibbs M.J., Jone R.A.C., 2017. The phylogenetics of the global population of potato virus Y and its necrogenic recombinants. Virus Evol. 3(1): vex002

Gouy, M., Guindon S., Gascuel O., 2010. SeaView version 4: a multiplatform graphical user interface for sequence alignment and phylogenetic tree building. Mol. Biol. Evol. 27, 221–224.

Gulya, T.J., Shiel, P.J., Freeman, T., Jordan, R.L., Isakeit, T., and Berger, P.H., 2002. Host range and characterization of sunflower mosaic virus. Phytopathology 92, 694–702.

Jones, R.A.C., 2009. Plant virus emergence and evolution: Origins, new encounter scenarios, factors driving emergence, effects of changing world conditions, and prospects for control. Virus Res. 141, 113–130.

Kaliciak, A., Syller J., 2009. New hosts of potato virus Y (PVY) among common wild plants in Europe. Eur. J. Plant Pathol. 124, 707–713.

Kiehr-Delhey, M., Delhey, R., 1981. Necochense mosaic of sunflower in the southeast of the province of Buenos Aires, Argentina. IV Jornadas Fitosanitarias Argentinas, UNC. 449–454.

Lemey, P., Rambaut, A., Drummond, A.J., Suchard, M.A., 2009. Bayesian phylogeography finds its roots. PLoS Comput. Biol. 5, e1000520.

Lemey, P., Rambaut A., Welch J.J., Suchard M.A., 2010. Phylogeography takes a relaxed random walk in continuous space and time. Mol. Biol. Evol. 27, 1877–1885.

Lenardon, S.L., 1994. Síntomas de etiología viral en cultivos de girasol. In: Pereyra, V.R., Escande, A.R. (eds.) Enfermedades del girasol en la Argentina. INTA, Balcarce, Buenos Aires. pp. 99–103.

Lenardon, S.L., Giolitti, F., León, A., Bazzalo, M.E., Grondona, M., 2001. Effects of Sunflower chlorotic mottle virus infection on sunflower yield parameters. Helia 24. 55–66.

Martin, D.P., Murell, B., Golden, M., Khoosal, A., Muhire, B., 2015. Rdp4: Detection and analysis of recombination patterns in virus genomes. Virus Evol. 1: vev003.

Minin, V.N., Bloomquist, E.W., Suchard, M.A., 2008. Smooth skyride through a rough skyline: Bayesian coalescent-based inference of population dynamics. Mol. Biol. Evol. 25, 1459–1471.

Muhire, B.M., Varsani, A., Martin, D.P., 2014. SDT□: A virus classification tool based on pairwise sequence alignment and identity calculation. PloS one, 9: e108277.

Patiño, V., 1963. Plantas introducidas. In: Plantas cultivadas y animales domésticos en América Equinoccial. Imprenta Departamental, Cali, Colombia. 542 p.

Rambaut, A., Drummond, A.J., Xie, D., Baele, G., Suchard, M.A., 2018. Posterior summarization in Bayesian Phylogenetics using Tracer 1.7. Syst. Biol. 67, 901–904.

Rieux, A., Khatchikian, C.E., 2017. TipDatingBeast: an R package to assist the implementation of phylogenetic tip-dating tests using BEAST. Mol. Ecol. Resour. 17, 608–613.

Ronquist, F., Teslenko, M., van der Mark, P., Ayres, D.L., Darling, A., Höhna, S., Larget, B., Liu, L., Suchard, M.A., Huelsenbeck, J.P., 2012. Mrbayes 3.2: Efficient bayesian phylogenetic inference and model choice across a large model space. Syst. Biol. 61, 539–542.

Simmons, H.E., Holmes, E.C., Stephenson, A.G., 2008. Rapid evolutionary dynamics of Zucchini yellow mosaic virus. J. Gen. Virol. 89, 1081–1085.

Sharman, M., Thomas, J.E., Persley, D.M., 2008. First report of Tobacco streak virus in sunflower (*Helianthus annuus*), cotton (*Gossypium hirsutum*), chickpea (*Cicer arietinum*) and mung bean (*Vigna radiata*) in Australia. Australas. Plant Dis. Notes 3, 27–29.

Suchard, M.A., Lemey, P., Baele, G., Ayres, D.L., Drummond, A.J., Rambaut, A., 2018. Bayesian phylogenetic and phylodynamic data integration using BEAST 1.10. Virus Evol. 4, vey016.

Xia, X., 2017. DAMBE6: new tools for microbial genomics, phylogenetics and molecular evolution. J. Hered. 108, 431–7.

Yasaka, R., Fukagawa, H., Ikematsu, M., Soda, H., Korkmaz, S., Golnaraghi, A., Katis N., Ho, S.Y.W., Gibbs, A.J., Ohshima, K., 2017. The timescale of emergence and spread of turnip mosaic potyvirus. Sci. Rep. 7(1), 4240.

