## Supplemental Table 1 for "Phylodynamics of sunflower chlorotic mottle virus, a re-emerging pathosystem"

**Table S1** Geographical localization, collection date, host and accession numbers of sunflower chlorotic mottle virus isolates infecting sunflower and weeds in Argentina.

| **Province** | **Department** | **Isolate-Location** | **Collection Date** | **Host** | **GenBank**  **Accession number** | **GPS data** | **Map^a^** |
| --- | --- | --- | --- | --- | --- | --- | --- |
| Entre Ríos | Paraná | Hasenkamp | Jan. 2008 | *H. annuus* | MG885791 | -31.526, -59.862 | 1 |
| Entre Ríos | Paraná | El Palenque | Jan. 2008 | *H. annuus* | MG885771 | -31.655, -60.186 | 2 |
| Entre Ríos | Paraná | Colonia Avellaneda | Jan. 2008 | *D. fullonum* | MG885788 | -31.773, -60.390 | 3 |
| Entre Ríos | Paraná | Oro Verde | Jan. 2008 | *H. annuus* | MG885770 | -31.848, -60.536 | 4 |
| Entre Ríos | Gualeguay | Gualeguay | Oct. 2011 | *D. fullonum* | MG885785 | -33.076, -59.399 | 5 |
| Córdoba | Marcos Juárez | Marcos Juárez | Apr. 2009 | *H. annuus* | MG885779 | -32.671, -62.065 | 6 |
| Córdoba | Gral San Martín | La Laguna | Oct. 2011 | *D. fullonum* | MG885786 | -32.727, -63.245 | 7 |
| Santa Fe | Gral López | Venado Tuerto | Oct. 2011 | *D. fullonum* | MG885782 | -33.705, -62.015 | 8 |
| Santa Fe | Gral López | Venado Tuerto | Oct. 2011 | *Eryngium* sp. | MG885787 | -33.705, -62.015 | 9 |
| La Pampa | Realicó | Realicó | Jan. 2017 | *D. fullonum* | MG885798 | -35.054, -64.209 | 10 |
| Buenos Aires | Pergamino | Pergamino | Oct. 2011 | *D. fullonum* | MG885781 | -33.916, -60.572 | 11 |
| Buenos Aires | Rojas | Rafael Obligado | Oct. 2011 | *D. fullonum* | MG885790 | -34.318, -60.745 | 12 |
| Buenos Aires | Gral Arenales | Gral Arenales | Jan. 2017 | *D. fullonum* | MG885793 | -34.317, -61.166 | 13 |
| Buenos Aires | 25 de mayo | Gobernador Ugarte | Feb. 2010 | *D. fullonum* | MG885784 | -35.118, -60.109 | 14 |
| Buenos Aires | Gral Alvear | Gral Alvear | Jan. 2017 | *D. fullonum* | MG885794 | -35.949, -59.870 | 15 |
| Buenos Aires | San Miguel  del Monte | San Miguel  del Monte | Oct. 2011 | *D. fullonum* | MG885783 | -35.474, -58.850 | 16 |
| Buenos Aires | Tandil | Tandil | Nov. 2006 | *D. fullonum* | MG885780 | -37.356, -59.204 | 17 |
| Buenos Aires | Tandil | Tandil | Nov. 2006 | *Ciconium* sp. | MG885792 | -37.219, -59.228 | 18 |
| Buenos Aires | Balcarce | Napaleofú | Jan. 2009 | *H. annuus* | MG885773 | -37.617, -58.746 | 19 |
| Buenos Aires | Lobería | La Alianza | Jan. 2009 | *H. annuus* | MG885774 | -37.711, -58.722 | 20 |
| Buenos Aires | Lobería | Dos Naciones | Jan. 2009 | *H. annuus* | MG885772 | -37.763, -58.652 | 21 |
| Buenos Aires | Balcarce | Los Pinos | Jan. 2009 | *D.fullonum* | MG885778 | -37.934, -58.345 | 22 |
| Buenos Aires | Gral Alvarado | Mechongué | Jan. 2009 | *H. annuus* | MG885777 | -38.136, -58.249 | 23 |
| Buenos Aires | Gral Alvarado | Miramar | Jan. 2009 | *H. annuus* | MG885776 | -38.269, -58.155 | 24 |
| Buenos Aires | Necochea | Quequén | Jan. 2009 | *H. annuus* | MG885789 | -38.535, -58.675 | 25 |
| Buenos Aires | Lobería | Pieres | Jan. 2017 | *H. annuus* | MG885795 | -38.389, -58.674 | 26 |
| **Province** | **Department** | **Isolate-Location** | **Collection Dates** | **Host** | **GenBank**  **Accession number** | **GPS data** | **Map^a^** |
| Buenos Aires | Lobería | Lobería | Jan. 2009 | *H. annuus* | MG885775 | -38.112, -58.740 | 27 |
| Buenos Aires | Lobería | Lobería | Jan. 2017 | *H. annuus* | MG885799 | -38.027, -58.725 | 28 |
| Buenos Aires | San Cayetano | San Cayetano | Jan. 2017 | *H. annuus* | MG885796 | -38.510, -59.695 | 29 |
| Buenos Aires | Tres Arroyos | Tres Arroyos | Jan. 2017 | *H. annuus* | MG885797 | -38.342, -60.406 | 30 |
| Entre Ríos | Paraná | Paraná | Jan. 2008 | *H. annuus* | GU181199^*^ | -31.846, -60.533 | 31 |
| Entre Ríos | Paraná | Colonia Avellaneda | Aug. 2008 | *I. lutea* | JN863232^*^ | -31.775, -60.378 | 32 |
| Buenos Aires | Lobería | Pieres | Jan. 2006 | *D.fullonum* | JN863233^*^ | -38.409, -58.668 | 33 |
| Buenos Aires | Lobería | Pieres | Jan. 2006 | *H. annuus* | GU181200^*^ | -38.409, -58.668 | 34 |

^*^: This sequences do not were obtained from this study; a: Map references used in Fig. 1.
