## Supplemental Table 2 for "Phylodynamics of sunflower chlorotic mottle virus, a re-emerging pathosystem"

**Table S2.** Bayes Factors (BF) for comparison of different diffusion models with Marginal likelihood estimators calculated using Stepping Stone (SS) method.

|  |  | **Log10 BF (model in row\|model in column)**^a^ | | | |
| --- | --- | --- | --- | --- | --- |
| **Model** | **ln MLE (SS)** | **Homogeneous (BD)** | **Gamma** | **Cauchy** | **LogNormal** |
| **Homogeneous (BD)** | -3741.18 | - | -11.27 | -9.98 | -3.77 |
| **Gamma** | -3715.24 | 11.27 | - | 1.28 | 7.49 |
| **Cauchy** | -3718.19 | 9.98 | -1.28 | - | 6.21 |
| **LogNormal** | -3732.49 | 3.77 | -7.49 | -6.21 | - |

^a^ BF should be interpreted as how many times the model in the row fits better to the data than the model in the column. Interpretation: log10(BF) > 1 is indicative of strong evidence, whereas log10(BF) > 2 is indicative of very strong evidence (Jeffreys, 1961).

**References**

Jeffreys, H., 1961. The Theory of Probability (3e). Oxford University Press, Oxford.
