## Supplemental Table 3 for "Phylodynamics of sunflower chlorotic mottle virus, a re-emerging pathosystem"

**Table S3.** Recombination events detected in the coat protein gene of sunflower chlorotic mottle virus isolates.

| **Coat protein gene** | | | | | | |
| --- | --- | --- | --- | --- | --- | --- |
| **Event /**  **Recombinant** | **Major parent** | **Minor parent** | **Breakpoint position** | | **Methods** | **P-value** |
|  |  |  | **begin** | **end** |  |  |
| 1/MG885794  General Alvear (BA) | MG885778  Los Pinos (BA) | GU181200  Pieres  (BA) | 482 | 773 | RDP | 7.425 x 10^-02^ |
|  |  |  |  |  | GENECONV | 7.050 x 10^-03^ |
|  |  |  |  |  | Bootscan | 1.686 x 10^-03^ |
|  |  |  |  |  | MaxChi | 2.286 x 10^-03^ |
|  |  |  |  |  | Chimaera | 1.567 x 10^-04^ |
|  |  |  |  |  | SiScan | 5.290 x 10^-07^ |
|  |  |  |  |  | 3Seq | 8.341 x 10^-04^ |

Recombination detection methods implemented in RDP4 software.
