## Supplementary figures and images for "Phylodynamics of sunflower chlorotic mottle virus, a re-emerging pathosystem"

### Supplemental Figure 1

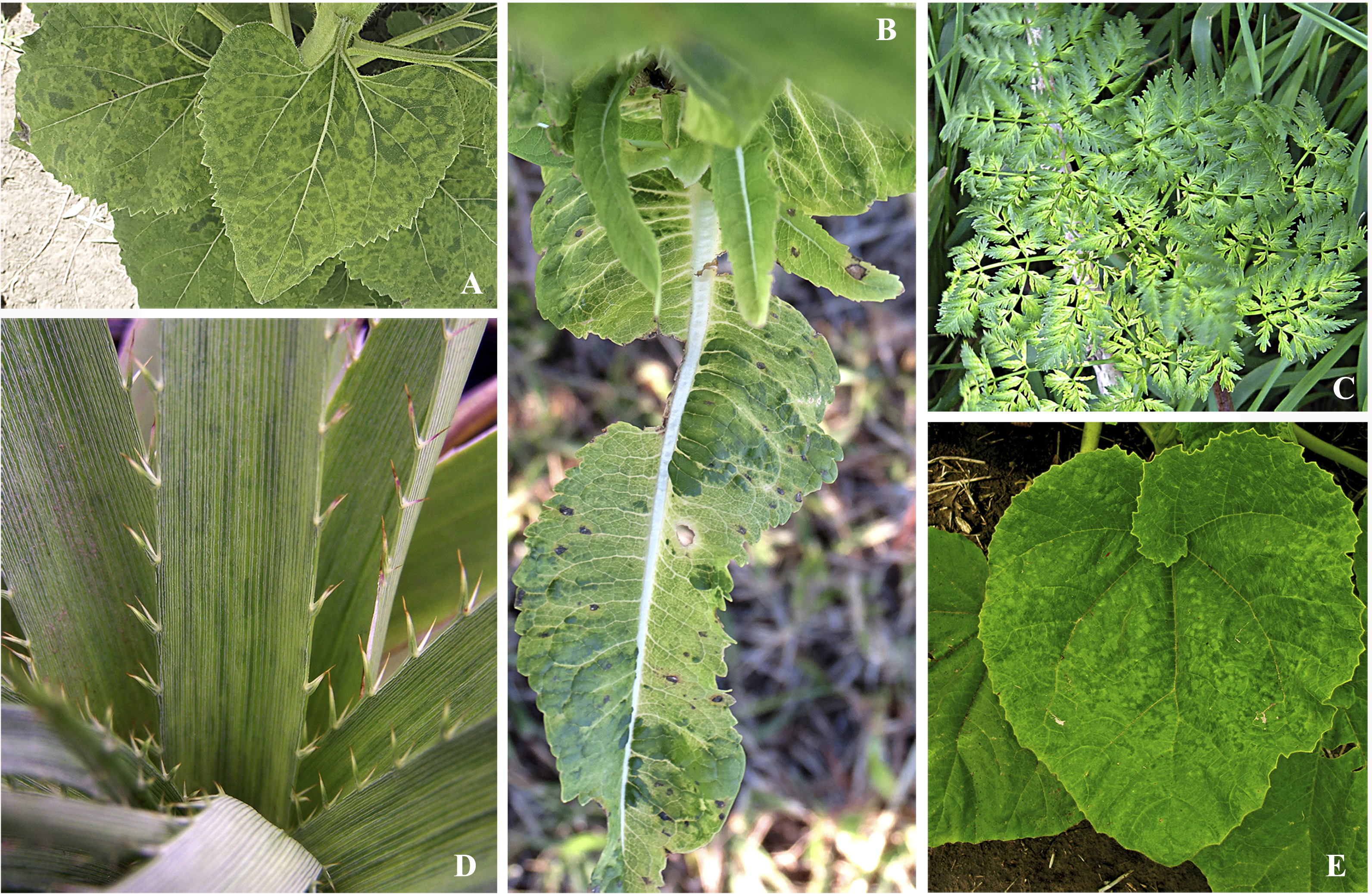

### Supplemental Figure 2

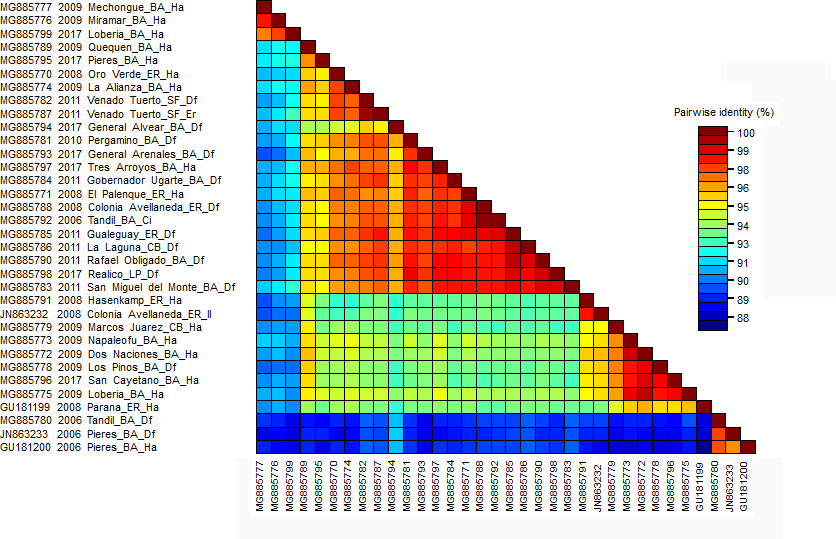
